# Divergent expression of *aristaless1* and *aristaless2* is associated with embryonic appendage and pupal wing development in butterflies

**DOI:** 10.1101/2022.09.28.509918

**Authors:** Erick X. Bayala, Isabella Cisneros, Darli Massardo, Nicholas W. VanKuren, Marcus R. Kronforst

## Abstract

Aristaless is a major regulator of developmental processes. It is well known for its role during appendage specification and extension across animals. Butterflies and moths have two copies of *aristaless, aristaless1* (*al1*) and *aristaless2* (*al2*), as a result of a gene duplication event. Previous work in *Heliconius* has shown that both copies appear to have novel functions related to wing color patterning. Here we expand our knowledge on the expression profiles associated with both ancestral and novel functions of Al1 across embryogenesis and wing pigmentation. Furthermore, we characterize Al2 expression, providing a comparative framework for understanding the role of gene duplicates in novel and ancestral roles. Our work shows that both Al1 and Al2 expression are associated with developing sensory appendages (leg, mouth, spines, and eyes) in embryos. Interestingly, Al1 appears to show higher expression earlier in embryogenesis while the highest levels of Al2 expression are shifted to later stages of embryonic development. Furthermore, Al1 localization appears extranuclear while Al2 co-localizes tightly with nuclei earlier, and then also expands outside the nucleus later in development. We observed similar cellular expression patterns for Al1 and Al2 in pupal wings when examining their roles in pigmentation. We also describe, for the first time, how Al1 localization appear to correlates with zones of Anterior/Posterior elongation of the body during embryonic growth, showcasing a possible new function related to Aristaless’ previously described role in appendage extension. Overall, these data suggest similar developmental roles associated with the extension/formation of specific appendages for both duplicates. However, we describe that such functions might be regulated by spatially and temporally complex patterns of expression for *al1* and *al2*. This work expands our knowledge of Aristaless function and expression following gene duplication and the implications of the duplication on butterfly development. Finally, and more fundamentally, our study helps clarify principles behind sub-functionalization and gene expression evolution associated with developmental functions following gene duplication events.

## Introduction

Across invertebrates, Aristaless has been show to function as a key regulator of proper patterning and appendage extension (Campbell et al., 1993; Schneitz, et al., 1993; Campbell & Tomlinson, 1998; Beermann and Schroder, 2004; Moczek, 2005). In flies, Aristaless was first described based on its role in the formation of a hair-like structure called the arista which extends from the antennae (Schneitz, et al., 1993). Since then, *aristaless* expression in flies has been shown to be relevant for the patterning of imaginal discs (both leg and wings) and their eventual extension into future adult structures (Schneitz, et al., 1993; Campbell & Tomlinson, 1998). Outside of flies, Aristaless has also been shown to play a role in the specification and extension of appendages. In *Gryllus*, early embryology work has shown how the expression of *aristaless* is associated with the tips of appendage buds growing out of the primary body plan (Miyawaki et al., 2002; Beermann and Schroder, 2004). Similarly, research in beetles has shown that the extension and branching patterns associated with their horns are also related to Aristaless activity (Moczek, 2005). Furthermore, there is evidence that *aristaless* plays a role in appendage formation in animals beyond insects. For example, velvet worms exhibit *aristaless* expression early in embryogenesis, within the cells forming leg buds. In cnidarians, the aristaless-like homebox gene (*alx*) is involved in initiating tentacle development during embryonic growth and across adult appendage regeneration (Smith et al., 2000) highlighting the presence of this function even before the protostome/deuterostome split. Even within echinoderms and vertebrates there is some evidence showcasing the role of Alx in skeletogenesis and limb patterning (Qu et al., 1998, Guerrero-Santoro et al., 2021). In summary, *aristaless* appears to be a key gene for the regulation of patterning, extension, and formation of appendages across several invertebrate species and perhaps even animals as a whole.

In Lepidoptera, the *aristaless* gene has been duplicated (Martin and Reed, 2010). Previous research on *aristaless* in butterflies and moths has mainly used a non-specific antibody that targets products of both copies of the gene (in addition to other homeodomain proteins; Martin and Reed, 2010). In moths, previous work has shown that one or both *aristaless* copies have important roles when it comes to the formation and proper patterning of the antennae (Ando, et al., 2018). Similarly, Aristaless activity in butterflies has been shown to be associated with color patterning processes during wing development (Martin and Reed, 2010). Furthermore, more specific approaches targeting their transcripts have suggested that different expression patterns for the two copies of the gene, *aristaless1* (*al1*), and *aristaless2* (*al2*), are involved in patterning specific color elements (Martin and Reed, 2010, Westerman et al., 2018), suggesting a novel role for pigmentation. Related to this novel color patterning function, our previous work studied *al1* in detail as the key regulator of the white and yellow color pattern switch in *Heliconius* butterflies (Westerman et al., 2018; Bayala et al., 2021). As part of that work, we showed that *al1* expression and activity are needed for the proper formation of embryonic appendages in *Heliconius* (Bayala et al., 2021). This developmental characterization suggested that Al1’s novel role in color patterning is possibly achieved by altering scale maturation or elongation rates, which functionally relates to the ancestral cellular processes of appendage formation and extension. The potential connection between Al’s functions related to color pattern formation and appendage formation underscores the need for a more thorough analysis of Al1’s role in appendage formation in *Heliconius*. This work would provide details on whether this role in appendage extension and pigmentation are rooted in the same cellular or developmental principles with respect to Al1 function.

Furthermore, no data are available related to Al2 expression or activity in the context of appendage formation or color patterning in *Heliconius*. Given there are two copies of the *aristaless* gene in Lepidoptera, proper understanding of how these genes evolved after the duplication event and/or potential regulatory or physical interactions requires that we characterize the developmental functions of both Al1 and Al2. Characterizing both genes in a comparative framework allows us to better understand the principles behind the developmental function of both Al1 and Al2 with respect to multiple aspects of *Heliconius* development. Furthermore, this approach provides an evolutionary angle, providing novel insight on how gene function can change following a duplication event.

Here we characterize Al1 and Al2 expression and developmental characteristics in a comparative framework across multiple aspects of butterfly development. We do this by first describing the expression pattern of Al1 across embryogenesis and in the wing across pupal development. We further extend this analysis to analyze Al2 expression and protein subcellular localization in order to gain a comparative view of both copies of the gene across *Heliconius* development. Finally, analyzing the similarities and differences between Al1 and Al2 expression and subcellular localization allows us to understand larger developmental principles behind their function in appendage formation and pigmentation.

To carry out this work we developed a series of tools for research in *Heliconius* butterflies. We used newly developed antibodies and *in situ* riboprobes that specifically target both gene products to tease apart expression differences between them. Furthermore, we applied these probes and antibodies to adapted protocols that allow us to analyze multiple embryonic stages for the first time in *Heliconius* butterflies. Embryos prior to hatching were used in order to analyze expression differences between Al1 and Al2 within appendage precursors and sensory organs. We coupled our early embryology analysis with pupal wing tissue staining to provide a complete view of the differences associated with Al1 and Al2 expression and subcellular localization across development and with respect to novel (wing coloration) and ancestral roles (appendage extension).

Our work provides evidence that expression of Al1 and Al2 is associated with sensory appendages during embryogenesis and pupal wing cells. More specifically, we show that Al1 activity in embryos is shifted earlier during appendage development and fades as development continues while Al2 expression is shifted to later stages of embryonic appendage development. A striking difference between Al1 and Al2 is observed in terms of subcellular localization because Al1 often appears more diffuse and extranuclear while Al2 appears to co-localize with nuclei. Furthermore, later in development, Al2 shows both nuclear and extranuclear expression. This temporal progression and these unique subcellular characteristics are also observed across pupal wing development, suggesting similar developmental functions between ancestral and novel roles. These observations suggest that both genes may still be involved with the process of appendage extension or formation, but they may be functioning in temporally distinct domains. Whether there is any physical interaction or compensation between them remains an open question. Our research provides the first description of the role of Al2 with respect to appendage extension in butterflies and compares it to our newly expanded information of Al1 and the ancestral single copy a*ristaless*. Finally, this work helps clarify how developmental profiles can shift following gene duplication events, possibly leading to divergent expression profiles and functional evolutionary differences.

## Results

### Expression of Al1 and Al2 is associated with the development of sensory appendages during mid (42-48 hours after deposition) embryogenesis

Our previous antibody staining showed that Al1 expression is associated with the development of appendages across specific stages of embryogenesis (Bayala et al., 2021). We used our newly developed Al1 probe to determine whether we could recapitulate this previously observed expression pattern for Al1 (Bayala et al., 2021). Both *in situs* and antibody staining showed that Al1 expression was associated with sensory appendages, exhibiting higher accumulation at the distal tip (**Figure 1 A’’-B**). This is consistent with our previous observations (Bayala et al., 2021) and with the expression of *aristaless* in other insects (Campbell et al., 1993; Schneitz, et al., 1993; Campbell & Tomlinson, 1998; Beermann and Schroder, 2004; Moczek, 2005). In addition, and as previously reported for Al1 (Bayala et al., 2021), the observed expression by antibody staining did not appear to co-localize with nuclei (**Figure 1A**).

**Figure 1:**
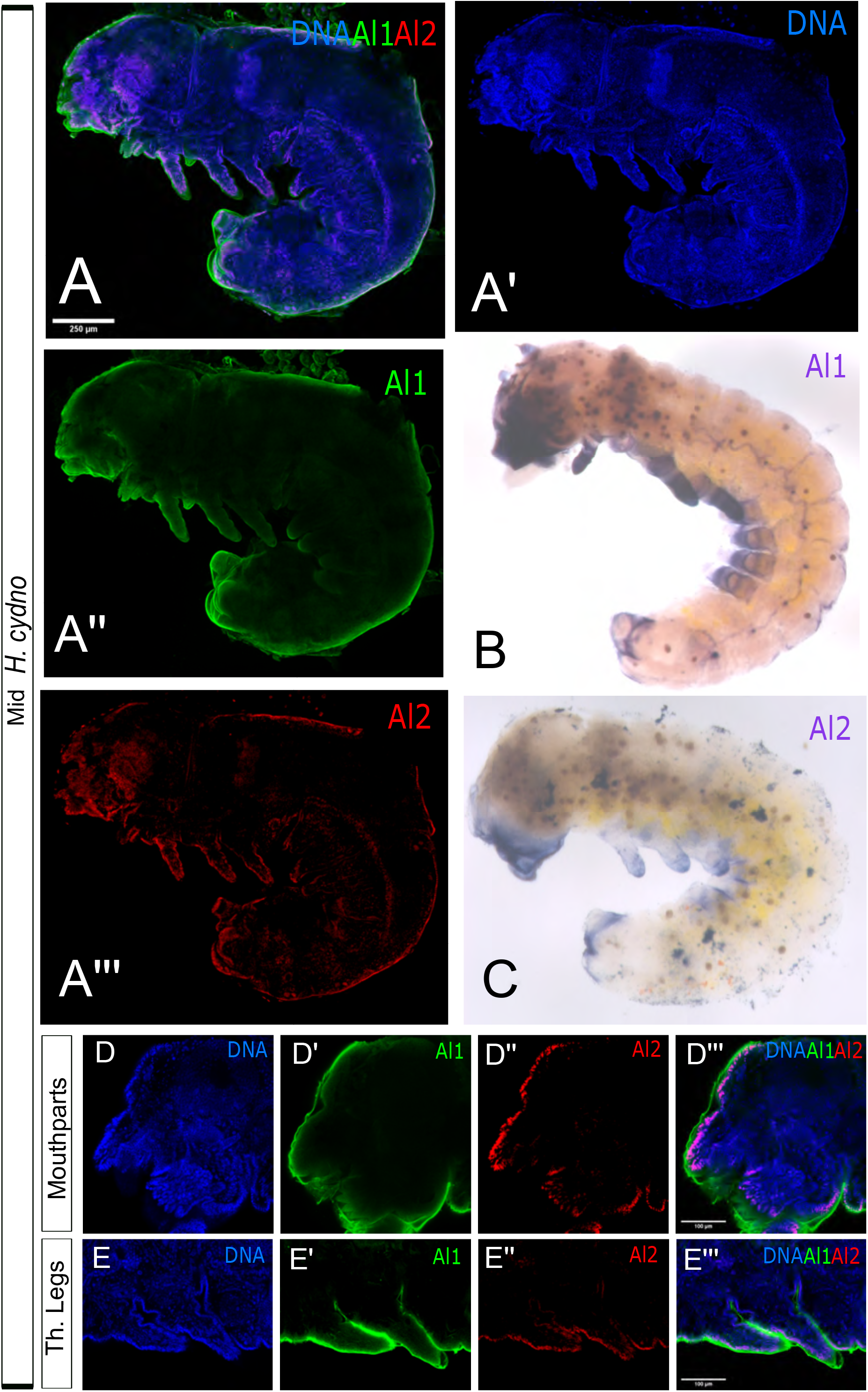
Immunodetection and in situ hybridization for Al1 and Al2 protein and transcript in mid (42-48 hours after deposition) wild-type *Heliconius cydno* embryos. (**A**) Immunodetection of Al1 and Al2 in a wild-type embryo (lateral-view). (**B**) Al1 *in situ* hybridization. (**C**) Al2 in situ hybridization. (**D-E**) Details on the Immunodetection of Al1 and Al2 in mouthparts (**D**) and thoracic legs (**E**) are shown. Panels show detection of DNA (A’, D, E), Al1 (A’’, D’, E’), Al2 (A’’’, D’’,E’’), and a merge (A, D’’’, E’’’). Scale bars are shown on the merge images.

We analyzed the same timepoint for Al2 and observed similar expression patterns between *in situ* and antibody staining across the same appendages (**Figure 1A’’’, C**). However, different from Al1, the expression of Al2 across appendages did co-localize with nuclei. Furthermore, Al2 expression appeared weaker than Al1 when comparing both within the same individual by antibody staining (**Figure A**) and when looking at two different embryos at the same developmental stage by *in situ* hybridization (**Figure 1B-C**). This suggests that both Al1 and Al2 are associated with the development of sensory appendages during *Heliconius* embryogenesis. However, the differences in subcellular localization within developing appendages and detection levels between both copies (**Figure 1D-E**) suggest other differences probably exist with respect to the expression pattern of both duplicates across development. To further characterize this, we analyzed earlier and later stages of embryonic development to get a clearer picture of how Al1 and Al2 expression changes across time.

### Al1 and Al2 expression patterns are temporally distinct and spatially complex but remain associated with embryonic sensory appendages across embryogenesis

We first analyzed embryos from earlier stages to determine if both Al1 and Al2 are expressed at the start of appendage growth. Our analysis of early embryos (36 to 42 hours after egg deposition) first revealed that Al2 was absent or very weakly expressed across the body and appendages while Al1 was highly expressed within developing sensory appendages (**Figure 2A**). Embryos at this time point showed strong expression of Al1 within multiple sensory appendages including mouth parts, thoracic legs, abdominal legs, eyes, and spines (**Figure 2A)**. The same pattern of expression for Al1 was observed by *in situ* hybridization (**Figure 2B**). As previously reported, the observed expression of Al1 did not appear to co-localize with nuclei (**Figure 2A, C**). Furthermore, CRISPR experiments supported the observed expression of Al1 at this time point because appendages lacking Al1 during this time point exhibited malformations and extension defects (**Figure 2D**).

**Figure 2:**
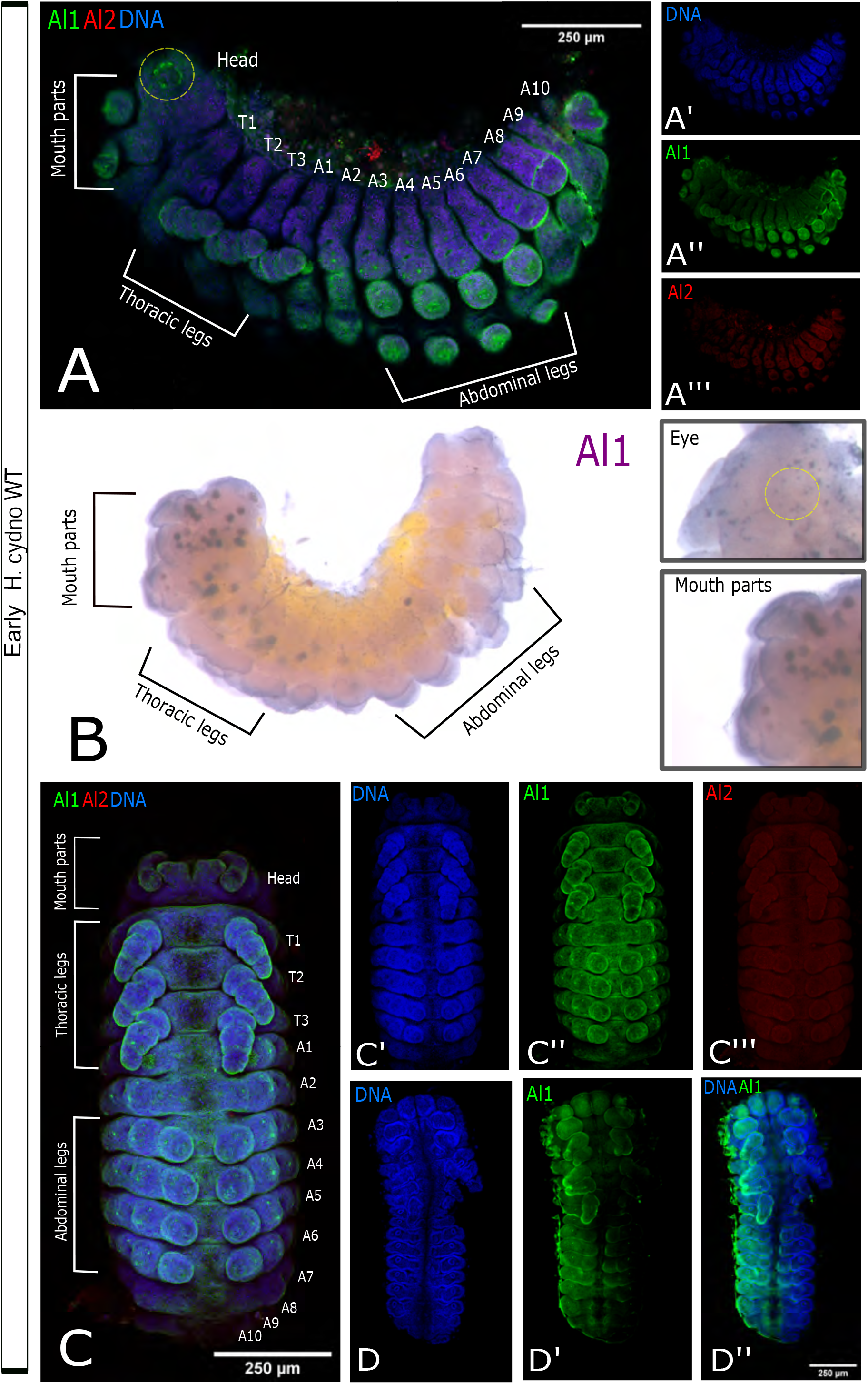
Immunodetection of Al1 and Al2 in early (36-42 hours after deposition) wild-type and –Al1 CRISPR *Heliconius cydno* embryos. (**A**) Immunodetection of Al1 and Al2 in a wild-type embryo (lateral-view). (**B**) Al1 *in situ* hybridization. (**C**) Immunodetection of Al1 and Al2 in a wild-type embryo (ventral-view). (**D**) Immunodetection of Al1 in an -Al1 CRISPR embryo (data from Bayala et al., 2022). White brackets highlight the different anatomical sections in A-B. The segments are also labeled on the merge images (Thoracic [T], Abdominal [A]). Panels show detection of DNA (A’, C’, D), Al1 (A’’, C’’, D’’), Al2 (A’’’, C’’’, D’’), and a merge (A, C, D’’). 250μm Scale bars are shown on all merge images.

We also analyzed and compared Al1 and Al2 expression patterns during late-embryonic development. During late time points (48-60 hours after egg deposition), we observed reduced Al1 expression across the entire embryo (**Figure 3 A-B**). Interestingly, during these late time points, and just a few hours before hatching, we detected strong Al2 expression both inside and outside of nuclei within multiple sensory appendages (mouth parts, thoracic legs, abdominal legs, eyes, and spines; **Figure 3A-G**). The high Al2 expression across these sensory appendages was also observed by *in situ* hybridization (**Figure 3C**). These observations across the majority of embryonic development showcase a temporal shift between the expression and possibly the activity of both copies of the gene. Such temporal shifts were also noticeable by *in situ* hybridization when looking across multiple stages of development using probes against both duplicates (**Supplemental Figure 1**). Furthermore, the differences we observed in terms of nuclear co-localization suggest different and dynamic spatial regulation between the duplicates.

**Figure 3:**
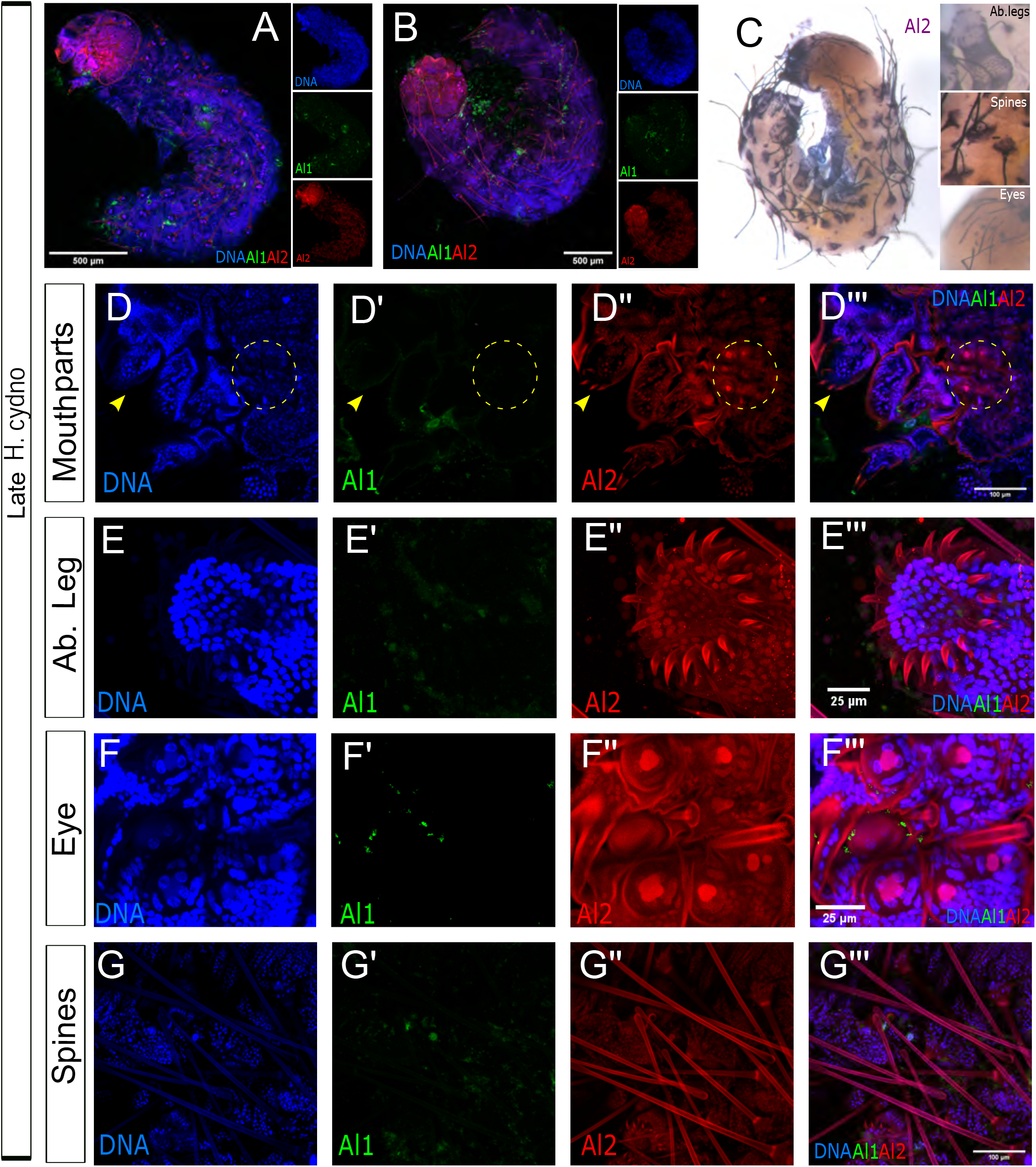
Immunodetection of Al1 and Al2 in early (48-60 hours after deposition) wild-type and –Al1 CRISPR *Heliconius cydno* embryos. (**A**) Immunodetection of Al1 and Al2 in a wild-type embryo (lateral-view). (**B**) Immunodetection of Al1 and Al2 in a wild-type embryo (dorsal-view). (**C**) Al2 *in situ* hybridization of wild-type embryo (**D-G**) Immunodetection of Al1 and Al2 in specific sensory appendages (mouthparts [**D**], Abdominal Leg tips [**E**], Eye [**F**], Spines [**G**]). Panels show detection of DNA (D, E, F,G), Al1 (C’, D’, E’, F’,G’), Al2 (C’’, D’’, E’’, F’’,G’’), and a merge (A, B, C’’’, D’’’, E’’’, F’’’,G’’’). Scale bars are shown on the merge images.

### Al1 and Al2 exhibit unique subcellular localization tightly associated with the development of spines and eyes

Our previous work described Al1’s role in appendage extension (Bayala et al., 2021). Most of that original description focused on embryonic legs and mouthparts. Here we further examined how expression patterns of Al1 and Al2 are associated with other sensory appendages (eyes and spines) across multiple stages of embryonic development. The development of these appendages has been poorly analyzed in terms of expression in other insects. Furthermore, analyses of Al1 and Al2 expression during the development of these sensory appendages do not exist in butterflies. We coupled our co-staining approaches with high magnification confocal microscopy to determine the expression patterns and subcellular localization of Al1 and Al2 during larval eye and spine development.

Similar to other appendages, eyes also exhibited strong Al1 expression early during embryonic development (**Figure 4A**) and strong Al2 expression later in development (**Figure 4B**). Early in eye development, Al1 was enriched in a ring-like pattern within the center of each one of the six simple eyes (ocelli) present on each side of the head. This ring-like pattern of expression was not co-localized with any of the nuclei in the center of the eye (**Figure 4A**). During the late stages of embryonic development, we observed Al2 in the center of the eyes, co-localizing with the three nuclei within each eye (**Figure 4B)**. Furthermore, there was clear detection of Al2 within a tube under each eye going deeper into the head/brain region (**Figure 4C**). These tube structures were visible within semi-transparent, freshly emerged caterpillars as a pigmented tube under the opening of the eyes (**Figure 4D**).

**Figure 4:**
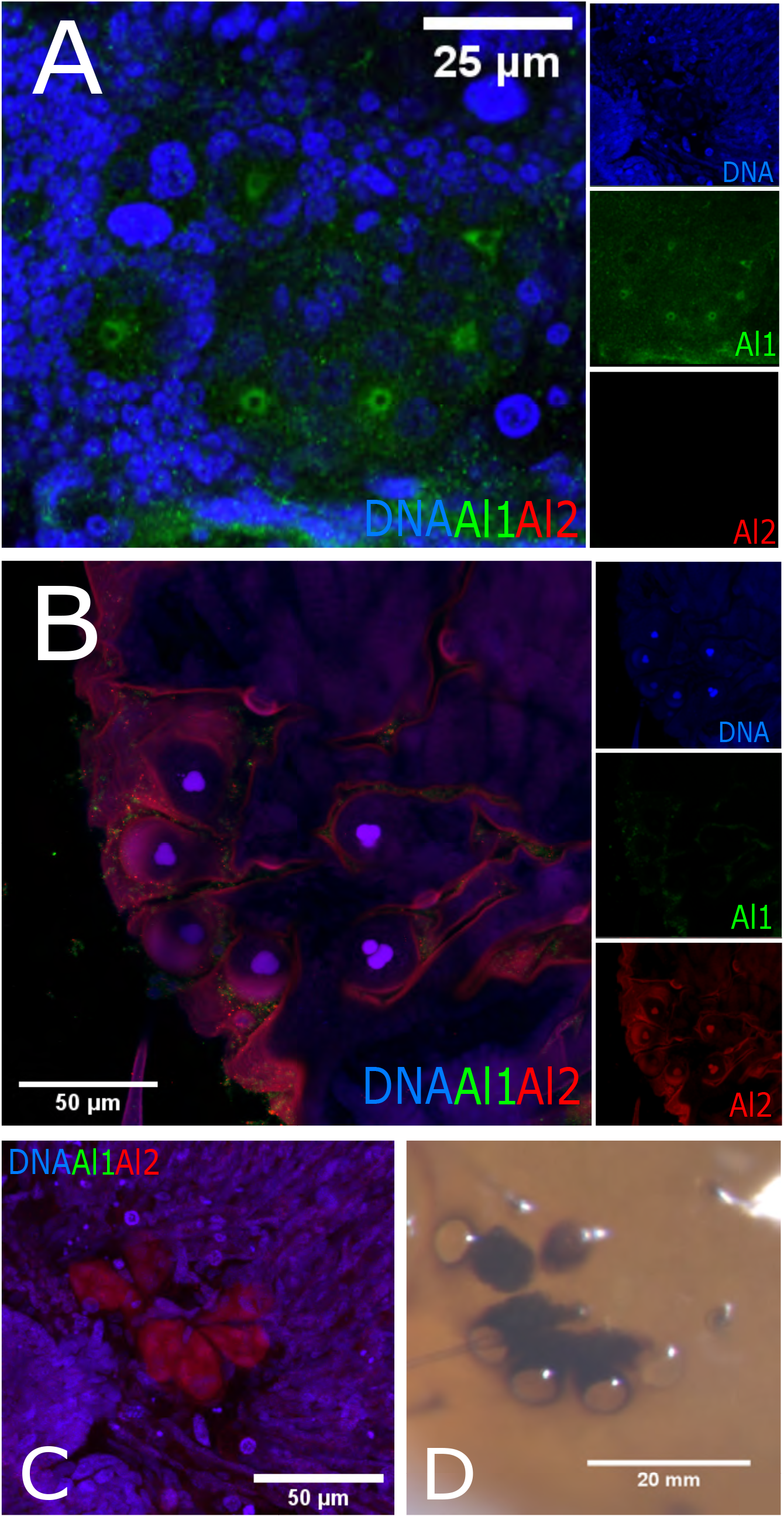
Detailed comparison of the Immunodetection of Al1 and Al2 between early (36 hours after deposition) and late (48 hours after deposition) wild-type eyes of *Heliconius cydno* embryos. (**A**) Immunodetection of Al1 and Al2 in early wild-type embryonic eyes. (**B**) Immunodetection of Al1 and Al2 in late wild-type embryonic eyes. (**C**) Deeper z-plane view of the B panel showcases detection of Al2 within the cell bodies projecting into the brain. (**D**) Caterpillar eye following eclosion. Panels show merge views (A, B, C). The insets of the merge images highlight DNA, Al1, and Al2 detection. Scale bars are shown on the merge images and Panel D.

We also analyzed spines during their development process. Like other sensory appendages, spines also exhibited a temporal shift between strong early Al1 expression (**Figure 5A**) and strong late Al2 expression (**Figure 5D)**. However, the bigger size of the spine cells and associated nuclei allowed us to observe several unique cellular features that we could not analyze in other appendages. As spine cells started to elongate, we observed the clear transition between Al1 and Al2 expression (**Figure 5A-D)**. Furthermore, it was very apparent that Al1 never co-localized with the two large nuclei that form the spine. Instead, Al1 appeared more diffuse along the entire spine body. Al2, on the other hand, was completely restricted to nuclei early in development (**Figure 5B**). Then, as development continued, a shift towards extranuclear Al2 was seen while still maintaining strong nuclear co-localization (**Figure 5C-D**). An intermediate state was observed as well where both Al1 and Al2 were detected within the developing spine (**Figure 5C**). Finally, during the late stages of development, one of the two nuclei that form part of the spine exhibited a higher level of Al2 than the other (the more distal nucleus within the spine exhibited higher Al2 expression; **Figure 5D**) possibly suggesting differences in the regulatory role of Al2 for different cell identities involved in forming the spine.

**Figure 5:**
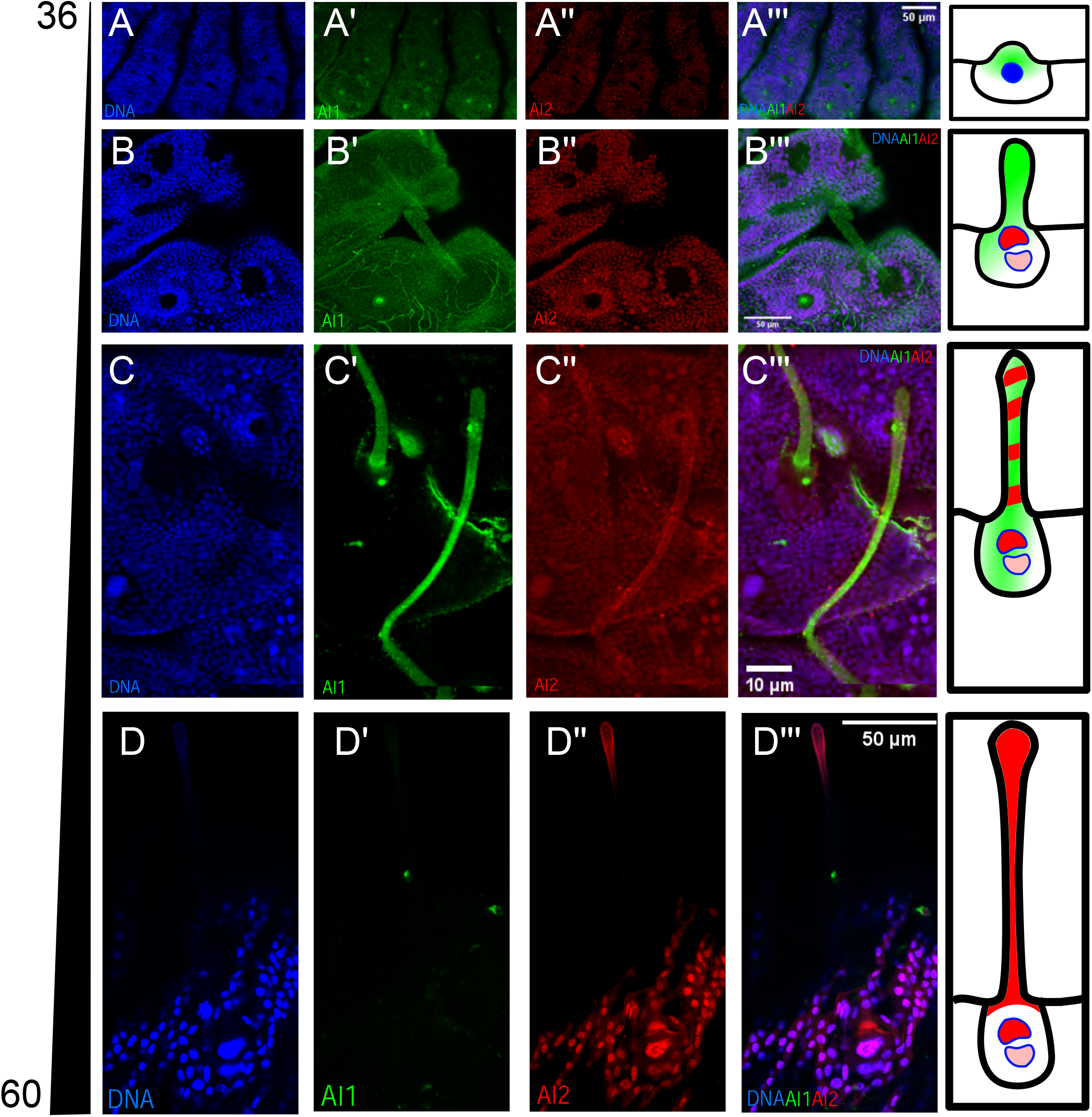
Detailed comparison of the Immunodetection of Al1 and Al2 across development in wild-type spines of *Heliconius cydno* embryos. (**A**) Immunodetection of Al1 and Al2 in wild-type embryonic spines 36 (**A**), 48 (**B**), 50 (**C**), and 60 (**D**) hours after deposition. Panels show detection of DNA (A, B, C, D), Al1 (A’, B’, C’, D’), Al2 (A’’, B’’, C’’, D’’), and a merge (A’’’, B’’’, C’’’, D’’’). Scale bars are shown on the merge images. A schematic representation of the observed subcellular detection for Al1 and Al2 is shown on the right of each time point.

### Al1 exhibits complex dorsal and ventral expression patterns in developing embryos

In addition to the described patterns of localization for Al1 within appendages, we also noticed an accumulation, either ventrally or dorsally, in developing embryos that has not been described for *aristaless* in other insects. When we analyzed Al1 across embryonic development we observed that Al1 protein expression correlated with parts of the embryos that were extending along the anterior-posterior (A/P) axis. Early in development, prior to the body fold that bends the legs inwards, we observed Al1 expression ventrally (**Figure 6A**). After the inward bend, and during A/P extension, we observed an accumulation of Al1 along the dorsal side of the embryo (**Figure 6B**). As A/P extension progressed, we observed a shift toward Al1 accumulation posteriorly (**Figure 6C**) coinciding with the extension of the abdomen. This localization along the dorsal side faded around 60 hours post egg deposition (**Figure 6D**) when the A/P extension is presumed to have concluded as the embryo is already occupying the entire physical space of the egg hours before hatching. The observed accumulation appeared more diffuse and did not coincide with nuclei, similar to what we observed in appendages.

**Figure 6:**
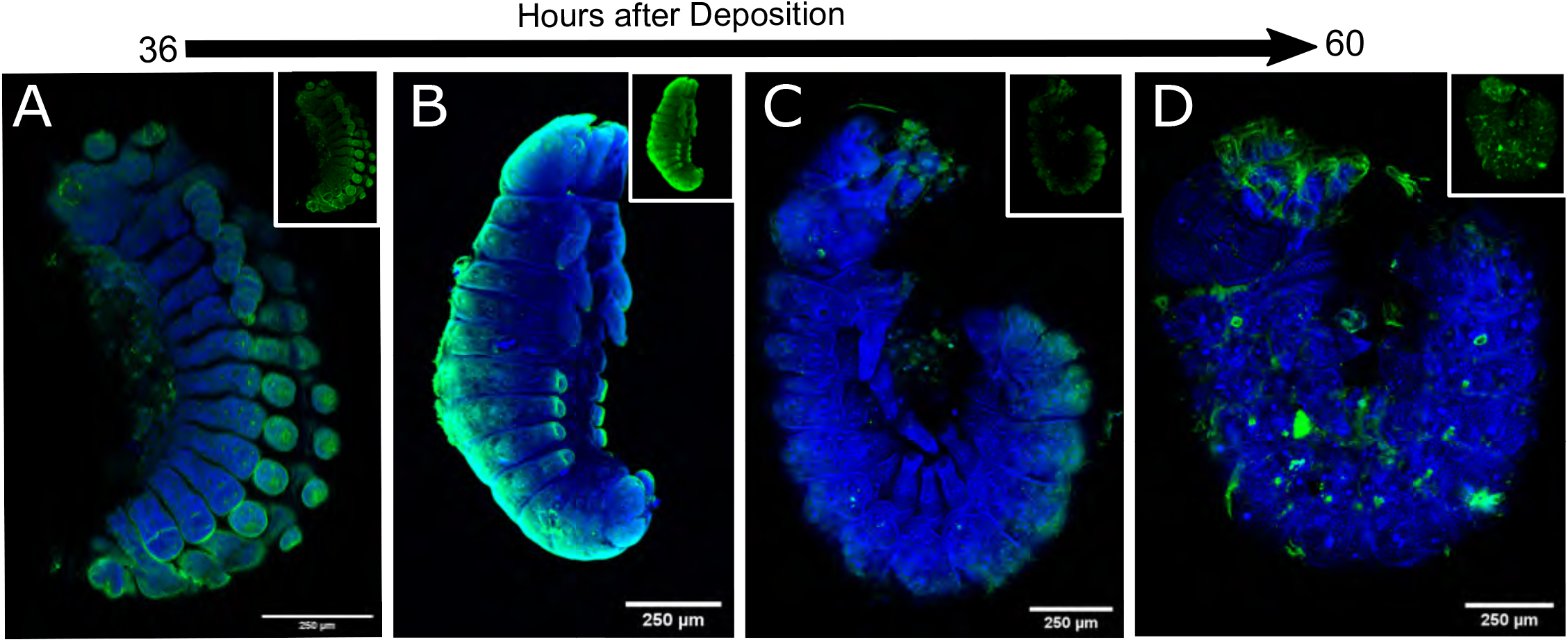
Immunodetection of Al1 across embryonic development in wild-type *Heliconius cydno* embryos highlighting detection of ventral and dorsal body sections extending within the anterior-posterior axis. Side-views are shown for the Immunodetection of Al1 in wild-type embryos at 36 (**A**), 42 (**B**), 50 (**C**), and 60 (**D**) hours after deposition. Panels show merged views of both DNA and Al1. Insets of just Al1 detection are also shown. Scale bars are shown on the merge images.

### Al1 and Al2 exhibit a temporal shift during wing development

The observed pattern of Al2 expression, transitioning from nuclear to extranuclear, and the temporal shift between early and late embryos raised the question about whether these observations are unique to the ancestral embryonic role of *aristaless* in appendage formation. We previously described how Al1 also exhibits a novel role with respect to color patterning in pupal wings (Bayala et al., 2021). Although, Al1 has been characterized developmentally as the switch between white/yellow coloration and an overall regulator of *Heliconius* wing coloration, such a novel role concerning pigmentation in *Heliconius cydno* has not been tested for Al2. However, some developmental data exist suggesting Al2 might also be involved in pigmentation (Martin and Reed 2010). Given the observed temporal transition from Al1 to Al2 in embryos, we examined whether this transition occurs in wings as well, perhaps suggesting a level of physical or regulatory interaction needed for proper function in the context of color pattern formation. This would also further provide evidence that the developmental mechanisms between ancestral and novel functions might be related, as previously suggested (Bayala et al., 2021).

We examined Al2 expression during pupal wing scale development (functional cellular units responsible for butterfly wing pigmentation) following previously described detection of Al1 early in pupal wing development (Bayala et al., 2021). When analyzing pupal scale cells, we observed a trend similar to what we described in embryos. In wings one day after pupal formation, we saw strong Al1 expression along the scale buds coming off from the wing blade (**Figure 7A**). At this stage, we noticed low levels of Al2 co-localizing with nuclei (**Figure 7A**). As development continued, we observed that around 3 days after pupal formation, Al1 was restricted within the developing scale cells extending outside of the wing blade while Al2 was still detected in the nuclei but also appeared around the cells within the wing blade (**Figure 7B-D**). Interestingly, Al2 appeared to be excluded from the nuclei of epidermal cells (**Figure 7B-D**) which are not believed to be involved in pigmentation (Nijhout, 1992). Around 4 days after pupal formation, we observed that Al2 still co-localized with nuclei but it also accumulated on the proximal parts of scale cells (**Figure 7E-G**). This was spatially distinct with Al1 which, as development continued, tended to localize more distally within scale cells (**Figure 7E-G**). Overall, the detection of Al2 appeared weaker than Al1 across the entire development process of the pupal wing, consistent with our previous measurements of gene expression (Westerman et al., 2018).

**Figure 7:** Immunodetection of Al1 and Al2 across pupal development in wild-type *Heliconius cydno*. Immunodetection of Al1 and Al2 is shown in wild-type pupal wings 1 (**A**), 3 (**B-D**), and 4 (**E-G**) days after pupal formation (APF). For 3 days APF a view of the scale (**B**) and nuclei (**C**) z-levels are shown as well as a side reconstruction of the entire scale cell body (**D**). Similarly, For 4 days APF a view of the scale (**E**) and nuclei (**F**) z-levels are shown as well as a side reconstruction of the entire scale cell body (**G**). Panels show detection of DNA (A, B, C, D’, E, F, G’), Al1 (A’, B’, C’, D’’, E’, F’, G’’), Al2 (A’’, B’’, C’’, D’’’, E’’, F’’, G’’’), and a merge (A’’’, B’’’, C’’’, D, E’’’, F’’’, G). Scale bars are shown on the merge images.

## Discussion

Our work presents the first characterization of Al2 expression in embryonic appendages, providing a unique point of contrast with the ancestral role of appendage development previously described for *aristaless* in other insects (Campbell et al., 1993; Schneitz, et al., 1993; Campbell & Tomlinson, 1998; Beermann and Schroder, 2004; Moczek, 2005) and Al1 in butterflies (Bayala et al., 2021). We performed this characterization in a comparative framework by further describing the Al1 pattern of expression across multiple stages of embryonic development and including an analysis of Al2 across the same stages (Summary in **Figure 8**). We first described how Al1 expression is higher earlier in development while Al2 expression increases later. Despite the temporal shift in their expression levels, both copies of the gene are still associated with growing sensory appendages. Previously, we showed that Al1 was expressed in embryonic leg and mouth structures (Bayala et al., 2021), and here we expanded on these observations by showing that Al2 is expressed in the same structures, but in different patterns. We also reported that Al1 and Al2 localization is associated with the eyes and spines across embryonic development and characterized specific cellular events associated with their subcellular and temporal dynamics. In addition, we showed that Al1 is localized ventrally and dorsally, matching body sections extending A/P during embryonic development. This expression pattern has not previously been described for *aristaless* but it appears to highlight regions of body axis extension and fades as that extension process ends (**Figure 6**). These data, although related with possible cellular roles of elongation and growth, which are important in appendage formation, may indicate an undescribed function for Al1 and Al2 in the context of A/P elongation.

**Figure 8:**
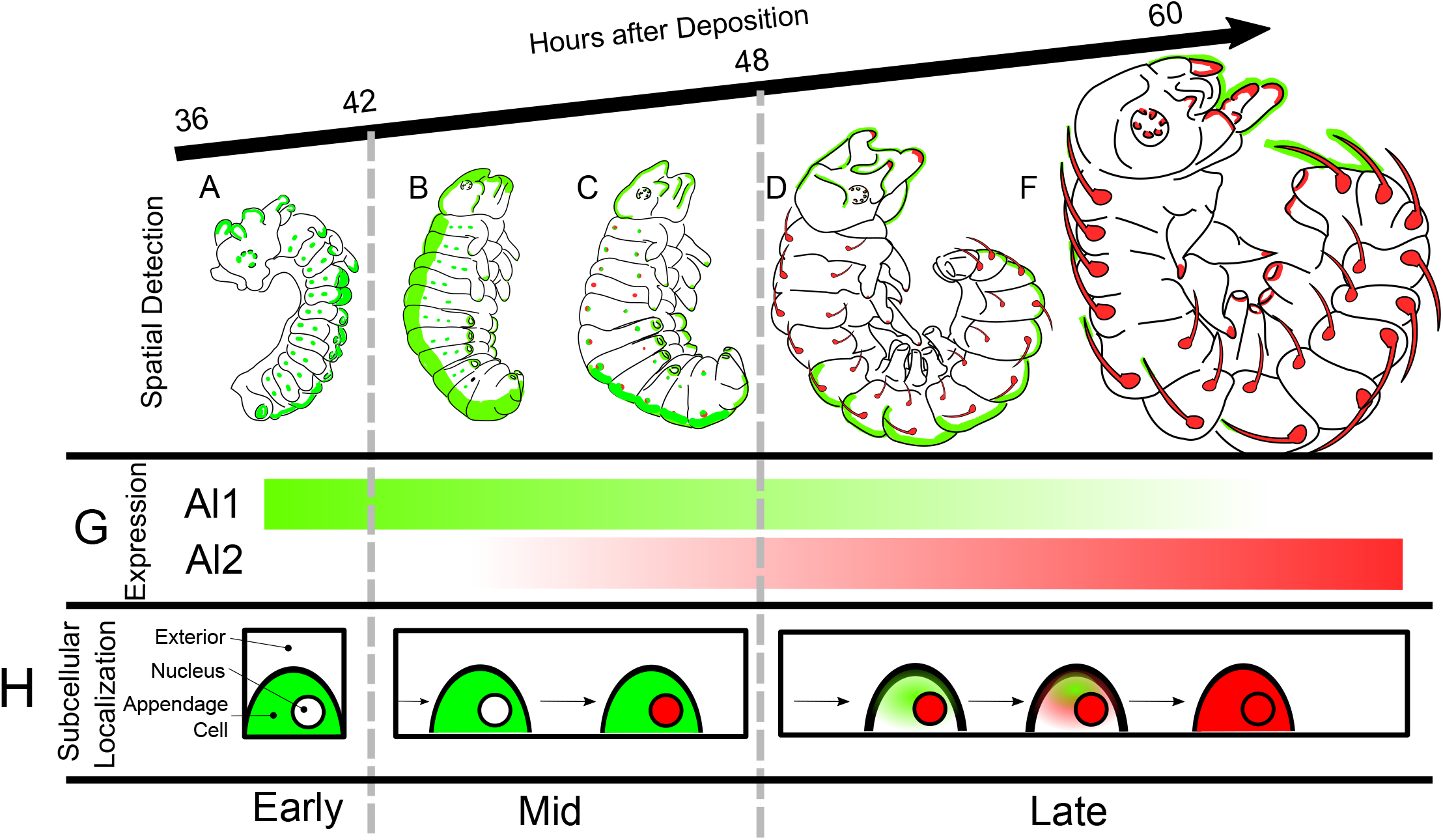
Summary of Al1 and Al2 Immunodetection in *Heliconius cydno* embryos across development. Schematics of the side-views of the embryos across development (between 36 to 60 hours after egg deposition) are shown. Al1 detection is shown in green while Al2 detection is shown in red within embryos in their early (**A**), mid (**B-C**), and late (**D-F**) developmental stages. (**G**) The bar graphs showcase the temporal shifts in the expression of Al1 and Al2. (**H**) Schematic representation of the observed subcellular localization for the detection of Al1 and Al2 over time during embryonic development.

Finally, we noted that Al2 exhibits co-localization with nuclei during mid and late stages of development. In addition, we also described how it exhibits a shift to extranuclear expression later in development. On the other hand, Al1 was always seen outside the nucleus (**Figure 8H)**, and this is consistent when using two different antibodies raised against two different Al1 peptides. These observations seem to be maintained even when looking at developing scale cell precursors on pupal wings, suggesting a possible underlying level of conservation with respect to their cellular function. Overall, our work describes some similarities between Al1 and Al2 with respect to their association with sensory appendages, possibly indicating similar upstream regulation. Some ancient paralogs, such as *engrailed* and *invected*, are known to be co-regulated during *Drosophila* development through the use of shared *cis-*regulatory elements (Cheng et al., 2014). Like *en/inv, al1/al2* were formed by tandem duplication and remain adjacent in the lepidopteran genome, perhaps suggesting a common mechanism for generating divergent expression patterns between nascent duplicates. However, we also note differences in terms of spatial and temporal control of their expression, perhaps suggesting differences at the functional level when it comes to their role in appendage formation and extension.

Our work characterizes expression differences between Al1 and Al2 within ancestral (sensory appendage formation) and novel roles (wing pigmentation) which, at its core, expands our views related to gene duplication, sub-functionalization, and the evolution of novel gene function (Long et al., 2013). It is well known that gene duplication can often lead to sub-functionalization by alleviating the responsibility imposed on a single copy (Force and Lynch, 1999). This process of sub-functionalization can be associated with shifts in spatial and temporal patterns of expression as a result of divergence in gene regulation between the two duplicates (Force and Lynch, 1999; Long et al., 2013). Following gene duplication and sub-functionalization, it is common to observe that the same overall gene function is maintained but now segmented into different parts of the developing body or across specific time windows as a result of divergent regulation (Force and Lynch, 1999). Neofunctionalization, on the other hand posits that one copy retains the ancestral function and the other gains a totally new function (Force and Lynch, 1999; Long et al., 2013). Furthermore, additional models exist of an in between state between these outcomes following gene duplication (Long et al., 2013).

Our data showcase an interesting, yet puzzling, case for the study of gene duplication and the evolution of ancestral and novel functions. First, we do not see spatially distinct expression domains in which the copies are restricted to specific appendages. However, we do see a shift in the temporal regulation of Al1 and Al2, suggesting that there has been at least some level of sub-functionalization at the temporal level associated with regulatory differences. However, the puzzling part resides in the strong differences observed in terms of subcellular localization. Al1 does not co-colocalize with nuclei (at least at the analyzed time points) while Al2 is observed to exhibit clear nuclear localization. Given that Aristaless has been described as a homeodomain transcription factor (Campbell & Tomlinson, 1998; Schneitz, et al., 1993) this could suggest that Al2 has maintained the function of transcriptional regulation. Another interesting observation is how the timing of Al2 activation matches the downregulation of Al1. These observations could suggest some antagonistic relationship between the two copies of the gene, as their expression patterns are temporally distinct with little overlap. This antagonistic relationship could provide the basis for novel mechanisms regulating appendage formation.

As we suggested previously (Bayala et al., 2021), it is possible that Al1 is regulating cellular processes outside the nucleus, as seen with other homeodomain proteins like Extradenticle (Abu-Shaar, et al., 1999). Furthermore, the entire subcellular dynamics we describe could be even more complex than expected. For example, it has been shown in vertebrates that Aristaless-Related Homeobox (ARX/ALX) genes can exhibit complex subcellular localization because of dimerization (homodimers and heterodimers) events and active sequestration via other transcription factors (Shoubridge et al., 2010; Guerrero et al., 2021). Future work should analyze the possibility that Al1 might be involved with the activation or repression of Al2. In both embryos and wings we observe a progression from Al1 to nuclear Al2, and given that in both embryos and wings knockouts Al1 produces a phenotype, this might imply that extranuclear Al1 activity is important for proper appendage formation or growth which, in part, could be mediated by Al2.

The similarities between the cellular events in embryos and wings might also suggest a similar underlying mechanism for both processes. It is known, for example, that Alx in rodents can cause pigmentation differences by affecting the maturation rate of pigment cells (Mallarino et al., 2016). Along with this idea, we believe similar parallels exist here between Al1 and Al2 regulating appendage growth and their possible role in scale maturation. Scale maturation has been shown to affect pigmentation outcomes via heterochronic shifts (Koch et al., 2000). So possible mechanisms involved in appendage growth might also be controlling scale extension and maturation. An experiment, where Al1 is knocked out and then Al2 is analyzed, could reveal the regulatory relationship between the two duplicates.

## Methods

### Butterflies rearing

Butterflies were reared in greenhouses at the University of Chicago with a 16h:8h light:dark cycle at ~27°C and 60% – 80% humidity. Adults were fed Bird’s Choice artificial butterfly nectar. Larvae were raised on *Passiflora oerstedii*.

### Embryos fixation and dissection

Eggs were collected from plants between 24 to 36 hours after deposition. I adapted the fixation scheme from Brakefield et al. (2009). Eggs were first transferred to 1.5 ml tubes and washed with PBS to remove any dirt. Eggs were then permeabilized and had their chorion removed with 5% Bleach (PBS) for 6 minutes. Eggs were then washed 5 times for 5 minutes in PBS to remove the excess bleach. We added 1 ml per tube of a 4% Paraformaldehyde solution (PBS) for fixing for 30 to 60 minutes. This fixation step was skipped for eggs being used for *in situ* hybridization and instead they were taken into the methanol series directly. Eggs exposed to Paraformaldehyde were then washed in PBST (PBS + 0.5% Triton-X100) 2 times for 5 minutes and then taken into a methanol series (25%, 50%, 75% methanol solutions in PBS at 4°C). Eggs were then transferred to 100% methanol and stored at -20°C for 5 days. Eggs were then transferred using plastic pipettes to a glass dissection plate with pre-chilled 100% methanol for dissection with fine forceps and dissection needles. Dissected embryos were then pipetted carefully into a 16 well tissue culture plate with 1 ml per well of chilled methanol. These embryos were taken back through a 1 ml per well methanol series (75%, 50%, 25% methanol solutions in PBS at 4°C) for rehydration in the case of antibody staining or maintained in 100% methanol for *in situ* staining. Then embryos were washed twice with 1 ml of PBST per well and stored in PBST at 4°C for antibody staining.

### Butterfly wing dissections

Butterflies were dissected at early pupal stages following Martin et al. (2014). The protocol and adaptations to it were carried out as follows. The pupae were anesthetized in ice for 20 mins before dissection. To obtain the pupal wings the pupae were pinned on the head and most posterior section of the body. The denticle belt was then removed using dissection forceps to allow for easier access to the wing. Then micro-dissection scissors were used to carefully cut around the wing margin using the pupal cuticle as a guide. The piece of cuticle together with the pupal forewing was removed and placed directly in a 16 well tissue culture plate with 1 ml per well of a 4% Paraformaldehyde solution for fixing. Pupal wings were fixed for 30 to 45 mins and then cleaned of any peripodial membrane by using fine forceps. After fixation, the tissue was then washed with PBST (PBS + 0.5% Triton-X100 for antibody staining five times to then be stored at 4°C until stained (not more than 30 days).

### *al1* and *al2 in situ* hybridization of *Heliconius* embryos

We designed and synthesized *al1* and *al2-*specific probes using the *H. cydno al1* and *al2* transcript model (selected region shows 100% identity with the target gene transcript model and around 60% identity with the other copy transcript model). 250 base-pair regions from *al1* and *al2* were amplified using primers (al1-forward GTTCCCTCGCAGCCATTCTT; al1-reverse TACGGCACTTCACCAGTTCT; al2-forward CACCTTTAACCCGACCTCCC; al2 reverse GCAGCTCGTGTTCTCTAGCA) by PCR, cloned into a TOPO vector (Invitrogen), and transformed into competent *E. coli* DH5a cells. We grew 3 replicates of 2 positive colonies and extracted DNA using a miniprep DNA extraction kit. We confirmed insert sequences via Sanger sequencing, linearized plasmids using Not1 and Sac1 restriction enzymes (New England Biolabs), and synthesized probes using a reverse transcription kit (Qiagen) with added DIG labeled nucleotides. The synthesized probes were purified using Qiagen RNAeasy columns.

*In situs* were performed following an adapted version of Ramos and Monteiro’s (2007) protocol designed for larval wings. The entire process was carried out in 24 well tissue culture plates. Tissue was stored in cold methanol post dissection and rehydrated into PBT (PBS1X 0.1%Tween20) the day of the experiment. Tissue was washed 5 times for 5 min with PBT, then incubated in a pre-hybridization buffer (50%formamide, 5XSSC, 0.1% Tween20, and 1mg/ml Salmon Sperm DNA) for 1 hour at 55°C. 1 ml of Hybridization buffer (50%formamide, 0.01g/ml glycine, 5XSSC, 0.1% Tween20, and 1mg/ml Salmon Sperm DNA) with approximately 50 ng of the target gene probe peer well and left to incubate at 55°C for at least 24 hours. The tissue was then washed 5 times for 5 min in pre-hybridization buffer and then left washing in pre-hybridization buffer for 24 hours at 55°C. Embryos were then blocked in 1% bovine serum albumin (BSA) in pre-hybridization buffer for 1 hr at 4°C. Anti-DIG antibody was added (1:2000) to each of the wells and incubated overnight at 4°C. The tissue was then washed with PBT extensively (10 times or more for 5 minutes) before development with BM-purple (1ml per well, Roche Diagnostics). Time of development was approximately 20 minutes at room temperature to 24 hours at 4°C depending on the probe. Stained tissue was imaged using Zeiss stereomicroscope Discovery.V20 with AxioCam adapter. Sense probes were used as controls for both duplicates.

### Al1 and Al2 antibody staining of embryos, larval, and pupal wings

We raised polyclonal antibodies against two Al1 peptides and 1 Al2 peptide using the company GenScript (New Jersey, USA). Peptide antigens (Al1-1: QSPASERPPPGSADC, Al1-2: DDSPRTTPELSHA, Al2: CGSGSGMDDEDIPRR) are located in the N-terminal 40 amino acids and share 25% and 30% identity between Al1 and Al2. Polyclonal antibodies were affinity purified after harvesting and tested for specificity by performing Dot blot tests as described.

We performed antibody staining in pupal wings following Martin et al. (2014). We also applied this staining protocol to embryos. Tissue stored in PBST (PBS, Tritonx) was blocked in 1% BSA in PBST for two hours, then incubated overnight in 1 mL blocking buffer and Al1 and/or Al2 specific antibodies (1:1000 for pupal wings and 1:3000 for embryos). Tissue was washed twice quickly, then 5 times for 5 mins in ~0.5 mL PBST, then incubated in 1 mL of the secondary staining solution (goat anti-rabbit-AlexaFluor 488 [Thermofisher] at 1:1000 for Al1 in pupae and 1:3000 for Al1 in embryos, Donkey anti-rat-AlexaFlour 555 [Thermofisher] at 1:1000 for Al2 in Pupae and 1:3000 for Al2 in embryos and Hoechst 33342 at 1:1000 [Thermofisher] in blocking buffer). The tissue was washed extensively and then mounted on glass slides using VectaShield (Vector Labs) on glass slides. Images were collected using a Zeiss LSM 710 Confocal Microscope and processed using Zen 2012 (Zeiss) and ImageJ. For wild-type Al1 and Al2 double antibody staining of embryos, we used and imaged about 5 individuals for all early, mid and late time points. For wild-type pupal wing stainings, we used forewings from 2 individuals by time point. For Al1 imaging across embryological development, we used a total of 5 embryos across different stages of development between 24 to 36 hours after egg deposition.

## Supplemental Figures

**Supplemental Figure 1:** *In situ* hybridization staining against *al1* and *al2* transcripts across *Heliconius* embryonic development. *In situ* hybridization staining are show in embryos spanning 36 to 60 hours after egg deposition for both Al1 (**A-D**) and Al2 (**E-H**). Multiple embryos of specific stages stained with control sense probes for both genes are also shown (**I-L**).

## References

1. Abu-Shaar, M., Ryoo, H. D., & Mann, R. S. (1999). Control of the nuclear localization of Extradenticle by competing nuclear import and export signals. Genes & Development, 13(8), 935. https://doi.org/10.1101/GAD.13.8.935

2. Ando, T., Fujiwara, H., & Kojima, T. (2018). The pivotal role of aristaless in development and evolution of diverse antennal morphologies in moths and butterflies. BMC Evolutionary Biology, 18(1), 1–12. https://doi.org/10.1186/S12862-018-1124-2/FIGURES/6

3. Bayala, E. X., VanKuren, N., Massardo, D., & Kronforst, M. (2021). From the formation of embryonic appendages to the color of wings: Conserved and novel roles of aristaless1 in butterfly development. BioRxiv, 2021.12.02.470931. https://doi.org/10.1101/2021.12.02.470931

4. Beermann, A., & Schröder, R. (2004). Functional stability of the aristaless gene in appendage tip formation during evolution. Development Genes and Evolution, 214(6), 303–308. https://doi.org/10.1007/S00427-004-0411-7

5. Brakefield, P. M., Beldade, P., & Zwaan, B. J. (2009). Immunohistochemistry staining of embryos from the African butterfly Bicyclus anynana. Cold Spring Harbor Protocols, 4(5). https://doi.org/10.1101/PDB.PROT5209

6. Brakefield, P. M., Beldade, P., & Zwaan, B. J. (2009). Fixation and Dissection of Embryos from the African Butterfly Bicyclus anynana. Cold Spring Harbor Protocols, 2009(5), pdb.prot5206. https://doi.org/10.1101/PDB.PROT5206

7. Campbell, G., Weaver, T., & Tomlinson, A. (1993). Axis specification in the developing Drosophila appendage: The role of wingless, decapentaplegic, and the homeobox gene aristaless. Cell, 74(6), 1113–1123. https://doi.org/10.1016/0092-8674(93)90732-6

8. Campbell, G., & Tomlinson, A. (1998). The roles of the homeobox genes aristaless and Distal-less in patterning the legs and wings of Drosophila. Development, 125(22), 4483–4493. https://doi.org/10.1242/DEV.125.22.4483

9. Carrel, J. E., & Nijhout, H. F. (1992). The Development and Evolution of Butterfly Wing Patterns. Annals of the Entomological Society of America, 85(6), 808–809. https://doi.org/10.1093/AESA/85.6.808

10. Cheng, Y., Brunner, A. L., Kremer, S., DeVido, S. K., Stefaniuk, C. M., & Kassis, J. A. (2014). Co-regulation of invected and engrailed by a complex array of regulatory sequences in Drosophila. Developmental Biology, 395(1), 131. https://doi.org/10.1016/J.YDBIO.2014.08.021

11. Force, A., Lynch, M., Pickett, F. B., Amores, A., Yan, Y. L., & Postlethwait, J. (1999). Preservation of duplicate genes by complementary, degenerative mutations. Genetics, 151(4), 1531. https://doi.org/10.1093/GENETICS/151.4.1531

12. Guerrero-Santoro, J., Khor, J. M., Açıkbaş, A. H., Jaynes, J. B., & Ettensohn, C. A. (2021). Analysis of the DNA-binding properties of Alx1, an evolutionarily conserved regulator of skeletogenesis in echinoderms. Journal of Biological Chemistry, 297(1). https://doi.org/10.1016/J.JBC.2021.100901

13. Koch, B. F., Lorenz, U., Brakefield, P. M., & ffrench-Constant, R. H. (2000). Butterfly wing pattern mutants: developmental heterochrony and co-ordinately regulated phenotypes. Dev Genes Evol, 210, 536-544.

14. Long, M., Vankuren, N. W., Chen, S., & Vibranovski, M. D. (2013). New Gene Evolution: Little Did We Know. Annual Review of Genetics, 47, 307. https://doi.org/10.1146/ANNUREV-GENET-111212-133301

15. Mallarino, R., Henegar, C., Mirasierra, M., Manceau, M., Scharadin, C., Vallejo, B. S., … Hoekstra, H. E. (2016). Developmental Mechanisms of Stripe Patterns in Rodents. Nature, 539, 518–523.

16. Martin, A., & Reed, R. D. (2010). Wingless and aristaless2 define a developmental ground plan for moth and butterfly wing pattern evolution. Molecular Biology and Evolution, 27(12), 2864–2878. https://doi.org/10.1093/MOLBEV/MSQ173

17. Martin, A., McCulloch, K. J., Patel, N. H., Briscoe, A. D., Gilbert, L. E., & Reed, R. D. (2014). Multiple recent co-options of Optix associated with novel traits in adaptive butterfly wing radiations. EvoDevo, 5(1), 1–14. https://doi.org/10.1186/2041-9139-5-7/FIGURES/6

18. Miyawaki, K., Inoue, Y., Mito, T., Fujimoto, T., Matsushima, K., Shinmyo, Y., Ohuchi, H., & Noji, S. (2002). Expression patterns of aristaless in developing appendages of Gryllus bimaculatus (cricket). Mechanisms of Development, 113(2), 181–184. https://doi.org/10.1016/S0925-4773(02)00020-5

19. Moczek, A. P., & Nagy, L. M. (2005). Diverse developmental mechanisms contribute to different levels of diversity in horned beetles. Evolution & Development, 7(3), 175–185. https://doi.org/10.1111/J.1525-142X.2005.05020.X

20. Qu, S., Tucker, S. C., Ehrlich, J. S., Levorse, J. M., Flaherty, L. A., Wisdom, R., & Vogt, T. F. (1998). Mutations in mouse Aristaless-like4 cause Strong’s luxoid polydactyly. Development, 125(14), 2711–2721. https://doi.org/10.1242/DEV.125.14.2711

21. Ramos, D., & Monteiro, A. (2007). In situ protocol for butterfly pupal wings using riboprobes. Journal of Visualized Experiments : JoVE, 4. https://doi.org/10.3791/208

22. Schneitz, K., Spielmann, P., & Noll, M. (1993). Erratum: Molecular genetics of aristaless, a prd-type homeo box gene involved in the morphogenesis of proximal and distal pattern elements in a subset of appendages in Drosophila (Genes and Development (1993) 7 (114-129)). Genes and Development, 7(5), 911. https://doi.org/10.1101/GAD.7.5.911

23. Shoubridge, C., Tan, M., Fullston, T., Cloosterman, D., Coman, D., McGillivray, G., Mancini, G., Kleefstra, T., & Gécz, J. (2010). Mutations in the nuclear localization sequence of the Aristaless related homeobox; sequestration of mutant ARX with IPO13 disrupts normal subcellular distribution of the transcription factor and retards cell division. PathoGenetics, 3(1), 1. https://doi.org/10.1186/1755-8417-3-1

24. Smith, K. M., Gee, L., & Bode, H. R. (2000). HyAlx, an aristaless-related gene, is involved in tentacle formation in hydra. Development, 127(22), 4743–4752. https://doi.org/10.1242/DEV.127.22.4743

25. Westerman, E. L., VanKuren, N. W., Massardo, D., Tenger-Trolander, A., Zhang, W., Hill, R. I., Perry, M., Bayala, E., Barr, K., Chamberlain, N., Douglas, T. E., Buerkle, N., Palmer, S. E., & Kronforst, M. R. (2018). Aristaless Controls Butterfly Wing Color Variation Used in Mimicry and Mate Choice. https://doi.org/10.1016/j.cub.2018.08.051

